# CRWN nuclear lamina components maintain the H3K27me3 landscape and promote successful reproduction in Arabidopsis

**DOI:** 10.1101/2023.10.03.560721

**Authors:** Junsik Choi, Mary Gehring

**Affiliations:** Whitehead Institute for Biomedical Research, Cambridge MA 02142; Dept. of Biology, Massachusetts Institute of Technology, Cambridge MA 02139

## Abstract

The nuclear lamina, a sub-nuclear protein matrix, maintains nuclear structure and genome function. Here, we investigate the role of Arabidopsis lamin analogs CROWDED NUCLEIs during gametophyte and seed development. We observed defects in *crwn* mutant seeds, including seed abortion and reduced germination rate. Quadruple *crwn* null genotypes were rarely transmitted through gametophytes. We focused on the *crwn1 crwn2* (*crwn1/2*) endosperm, which exhibited enlarged chalazal cysts and increased expression of stress-related genes and the MADS-box transcription factor *PHERES1* and its targets. Previously, it was shown that *PHERES1* is regulated by H3K27me3 and that CRWN1 interacts with the PRC2 interactor PWO1. Thus, we tested whether *crwn1/2* alters H3K27me3 patterns. We observed a mild loss of H3K27me3 at several hundred loci, which differed between endosperm and leaves. These data indicate that CRWNs are necessary to maintain the H3K27me3 landscape, with tissue-specific chromatin and transcriptional consequences.

## Introduction

Eukaryotic genomes are surrounded by nuclear envelopes, which interact with and can impact the function of the underlying genome^1^. The nuclear envelope is composed of outer and inner nuclear membranes, which are studded with nuclear pore complexes and nuclear membrane proteins^2^. In metazoan cells, the nucleoplasmic side of the inner nuclear membrane is covered by an interlaced protein layer called the nuclear lamina, which serves as a key platform for genome regulation due to its proximity to the DNA^3^. The nuclear lamina is essential and primarily composed of lamins in animals. Although plants do not have orthologous lamins, functional analogs called NMCP (Nuclear Matrix Constituent Protein) proteins, which were initially discovered by analyzing the nuclear matrix structure in carrots^4,5^, are found in land plants^6,7^. NMCPs have a tripartite structure that structurally resembles animal lamins^4,8–10^ and the coiled-coiled domain can form filamentous structures^8^. NMCPs have been most intensively studied in *Arabidopsis thaliana*, which has four NMCP orthologs called CRWN (CROWDED NUCLEI), or formerly known as LINC (LITTLE NUCLEI) proteins^11,12^. Mutations of *CRWN* genes alter nuclear morphology^11,12^, reminiscent of abnormal nuclear shape seen in human nuclear *LMNA* (Lamin A/C) gene mutants.

Metazoan lamins interact with both other proteins and chromatin^2^, which are referred to as lamina-associated proteins (LAPs) and lamina-associated domains (LADs), respectively. In Arabidopsis, protein-protein interactors of CRWNs were first focused on nuclear envelope proteins such as SUN1^13^ and KAKU4^14^. Recently, a growing number of other interactors have been reported, including the PRC2 complex interactor PWO1^15,16^. This suggests a possible epigenetic role for CRWN1, but epigenomic histone modification data has not been reported for *crwn* mutants. CRWN1 proteins also interact with regions of the genome known as plant lamina-associated domains (PLADs), which have been described as transcriptionally inactive and marked with repressive chromatin modifications^17^. In line with characteristics of PLADs, mutations in *crwn* genes, particularly in combination, cause mis-regulation of a significant number of genes in vegetative tissue^18–21^. However, the effect of *crwn* mutations in reproduction is relatively less studied, especially on the genome-wide scale.

Although NMCPs and CRWNs share some functional similarities with their animal lamin counterparts, mutations in Arabidopsis *CRWN* genes revealed phenotypes distinct from animal laminopathies. One such phenotype is spontaneous immune responses in *crwn* mutants^18–21^. *crwn1 crwn2* (*crwn1/2*) and *crwn1 crwn4* (*crwn1/4*) mutants have high levels of salicylic acid (SA), which are responsible for the observed diminished size of rosette leaves and the autonomous defense response^19^. Depletion of SA, however, does not fully suppress other phenotypes observed in *crwn* mutants, including silique and seed phenotypes^19^. Here, we further investigated the basis of reproductive phenotypes in gametophytes and seeds of *crwn* mutants. Seeds are comprised of the embryo, endosperm, and ovule integument/seed coat tissues. Among these, the endosperm is an epigenetically divergent tissue^22,23^ where CRWNs’ functions have not been investigated. We show that CRWNs are required for Arabidopsis gametophyte viability and contribute to H3K27me3 patterning as well as proper gene expression in both endosperm and leaves. Our transcriptomic and epigenomic data indicate that CRWNs are important for development and epigenetic states of each tissue while maintaining distinct genome-wide profiles between endosperm and leaves, showing that CRWNs roles are vital and can be tissue-specific.

## Results

### *crwn* mutants display gametophyte and seed phenotypes

We observed that plants with various combinations of *crwn* mutations have shorter siliques than WT (Fig. 1a,b). The length of *crwn1/2* double mutant siliques is approximately two-thirds of WT (Fig. 1b). In line with aberrant seed morphologies previously reported^19^, combining a mutation in the SA biosynthesis gene *SID2/ICS1* (*SALICYLIC ACID INDUCTION DEFICIENT 2*/*ISOCHORISMATE SYNTHASE1*) with *crwn1/2* did not rescue the reduced silique length of *crwn1/2* mutants (Extended Data Fig. 1a), suggesting that this phenotype is likley SA-independent. Silique length was further reduced in the triple mutant combinations *crwn1 crwn2 crwn4* (*crwn1/2/4*) and *crwn1 crwn3 crwn4* (*crwn1/3/4*) (Fig. 1a,b).

**Fig. 1.**
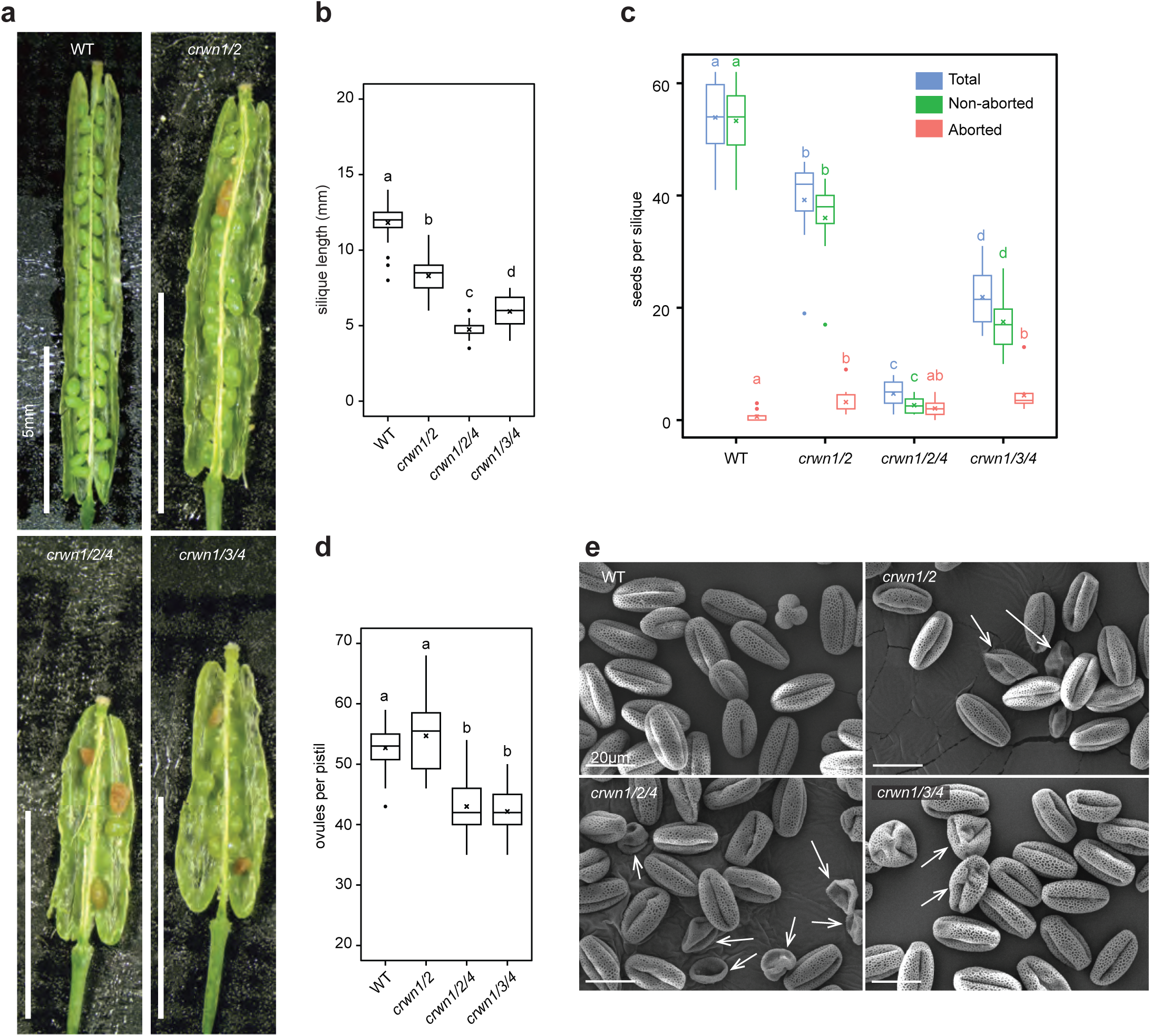
Reproductive phenotypes of *crwn* mutants. **a,** Representative silique images of WT, *crwn1/2*, *crwn1/2/4*, and *crwn1/3/4* mutants. Prematurely browning seeds with wrinkled surfaces are considered aborted Scale bars, 5 mm. **b,** Silique length of WT (*n* = 30), *crwn1/2* (*n* = 53), *crwn1/2/4* (*n* = 47), and *crwn1/3/4* (*n* = 34). Siliques were measured when they started turning yellow to ensure that growth was completed. **c,** Number of aborted and non-aborted seeds produced in siliques of WT (*n* = 10), *crwn1/2* (*n* = 10), *crwn1/2/4* (*n* = 14), and *crwn1/3/4* (*n* = 10). The total number of seeds is the sum of aborted and non-aborted seeds. **d,** Number of ovules in pistils of WT (*n* = 24), *crwn1/2* (*n* = 26), *crwn1/2/4* (*n* = 23), and *crwn1/3/4* (*n* = 17). Open flowers used to count ovules were from primary inflorescences of non-pruned plants. **e,** SEM images of pollen from each indicated genotype. Arrows indicate morphologically defective pollen. Scale bars, 20 µm. (**b-d**) Boxplots show median (centerline), upper and lower quartile (Q3 and Q1. box), Q1 - 1.5*interquartile range (IQR) and Q3 + 1.5*IQR (whiskers), outliers (points), and mean (‘x’ mark). Statistical analyses are one-way ANOVA with post hoc Tukey’s tests. For **c**, tests were performed among genotypes for the same seed type (the same color). Different letters indicate significant differences among them (*p* < 0.05).

Consistent with the shorter silique length, *crwn* mutants produced fewer seeds per silique, ranging from as few as an average 4.7 ± 2.2 seeds per silique in *crwn1/2/4* to 39.2 ± 8.2 seeds in *crwn1/2*, compared to 53.9 ± 6.9 seeds in the WT (Fig. 1c). The reduced seed set was only partially explained by reduced ovule number, with approximately 43.0 ± 4.6 ovules per pistil in *crwn 1/2/4* compared to 52.7 ± 3.8 ovules per pistil in the WT (Fig. 1d). The *crwn 1/2* mutant had a similar average number (54.7 ± 5.7) of ovules per pistil as the WT, suggesting that reduced ovule number was not a cause of reduced seed set in *crwn 1/2* (Fig. 1c,d). These data suggest that a high proportion of ovules remain unfertilized or have defects in post-fertilization development in *crwn* mutants. To investigate whether there are male gametophyte defects that might lead to reduced fertility, we investigated pollen morphology by scanning electron microscopy (SEM). We consistently observed abnormal, wrinkled, and collapsed pollen grains in all *crwn* mutant combinations (Fig. 1e). Thus, defective pollen could further contribute to reduced seed set. Several nuclear envelope protein mutants are known to produce pollen where the order of sperm cell nuclei (SN) and the vegetative cell nucleus (VN) in the pollen tube is altered compared to the WT^24,25^. We tested if *crwn* mutants suffer similar phenotypes. Although we observed a small increase of abnormal nuclear migration pattern in *crwn* pollen tubes, the degree was not significantly altered from the WT, which is similar to *crwn1* mutants tested by Goto et al^24^ (Extended Data Fig. 1b).

All *crwn* siliques showed evidence of seed abortion, which we quantified in each mutant combination (Fig. 1a,c). *crwn 1/2/4* exhibited the most extensive seed abortion, producing as many aborted seeds as normal seeds (Fig. 1a,c). Seeds from *crwn* mutants were also of variable size, shape, and appearance (Fig. 2a). To determine whether seed size variation had phenotypic consequences, we sorted *crwn1/2* seeds into two groups using a mesh with a 0.2328 mm opening. A fraction of both larger and smaller *crwn1/2* seeds were misshapen with an irregular, wrinkled surface (Fig. 2a). We then determined the germination rate of the two populations and the WT (Fig. 2b). The maximal germination of smaller *crwn1/2* mutant seeds, which passed through the mesh, was 33.1% at 10 days after stratification. Larger *crwn1/2* mutant seeds, those retained by the mesh, germinated at a higher rate than the small *crwn 1/2* seeds but at a lower rate than the WT – 90.2% vs 100% at 10 days.

**Fig. 2.**
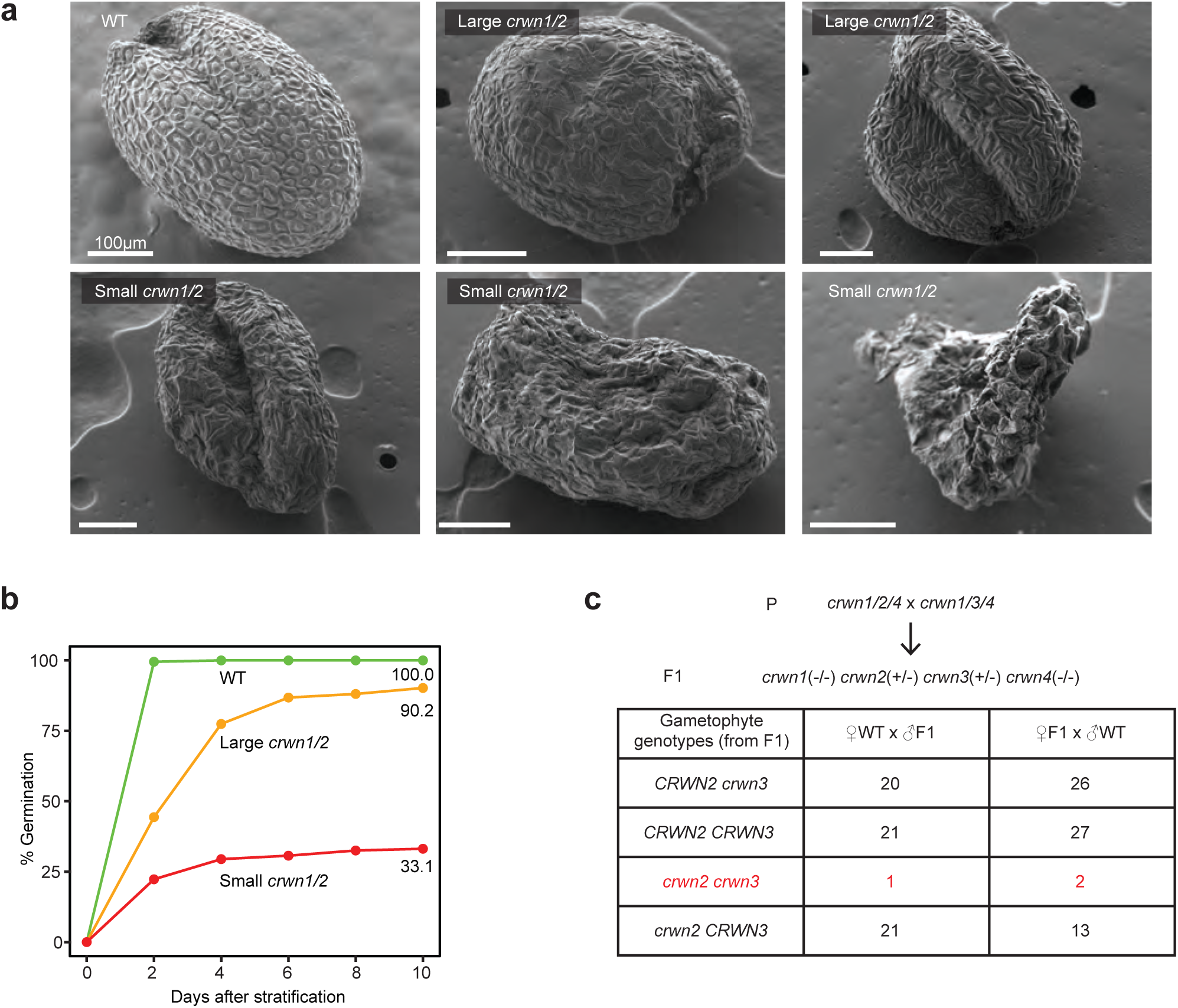
Loss of *CRWN*s leads to defective seeds and reduced gameto-phyte viability. **a,** SEM images of WT and *crwn1/2* seeds. Small and large seeds were defined depending whether they could pass through a mesh with 0.2328 mm opening. Scale bars, 100 µm. **b,** Germination rates of WT (*n* = 210), large *crwn1/2* seeds (*n* = 248), and small *crwn1/2* seeds (*n* = 157). **c,** Reciprocal crosses between WT and *crwn* mutant F1 plants. All gameto-phytes produced from F1 plants were mutant for *crwn1* and *crwn4* and either mutant or wild type for *crwn2* and *crwn3*. Red indicates quadruple *crwn* mutant gametophytes generated from F1 plants. Chi-squared test values for expected ratio of 1:1:1:1 are 18.46 (*p* < 0.005) for WT x F1 progeny and 24.8 (*p* < 0.005) for F1 x WT progeny.

It was previously shown that the quadruple *crwn* mutant could not be recovered^12^. However, it was unknown at which developmental stage this lethality occurred. We sought to assess the requirement of *CRWNs* for reproduction by determining transmission of the full complement of *crwn* mutations through the male and female gametophytes (Fig. 2c). We first crossed *crwn1/2/4* mutants with *crwn1/3/4* mutants to create F1 progeny that were homozygous for *crwn1* and *crwn4* mutations and heterozygous for *crwn2* and *crwn3*. We reciprocally crossed F1 plants to the WT and determined the genotypes of F2 plants. If *crwn1/2/3/4* is gametophytic lethal, we would not recover any F2 plants that are heterozygous for all four *CRWN* loci. (Note that although *CRWN2* and *CRWN3* are both on chromosome 1, they are approximately 80 cM apart and thus not genetically linked.) When WT females were pollinated with F1 males, we recovered one of 63 F2 plants that was heterozygous for all *CRWN* loci (Fig. 2c). This is significantly different from the expected 1:1:1:1 ratio of genotypes (χ^2^ = 18.46; *p* < 0.005). In the reciprocal cross, two heterozygous F2 individuals out of 68 were recovered (χ^2^ = 24.8; *p* < 0.005) (Fig. 2c). In this cross, there was also a reduction in the number of progeny whose female gametophytes were *crwn1 crwn2 CRWN3 crwn4*, suggesting additional sensitivity to *crwn2* mutation in the female gametophyte (Fig. 2c). This partially explains why *crwn1/2/4* has the lowest seed set of the observed genotypes (Fig. 1a,c). We conclude that at least one *CRWN* gene is required for gametophytic viability, with rare exceptions.

In summary, we identified a series of phenotypic defects in *crwn* mutant ovules, gametophytes, and seed. These mutant phenotypes vary in expressivity and penetrance, suggesting interaction between *CRWNs* and additional non-genetic factors.

### *crwn1/2* mutants exhibit enlarged endosperm chalazal cysts and variability in the timing of seed development

We focused the remainder of our investigations on *crwn1/2* because it has less severe phenotypes than *crwn* triple mutants and produces a sufficient number of seeds to facilitate experimentation. Because *crwn1/2* seeds exhibit morphological defects at maturity and a lower germination rate (Fig. 2a,b), we more closely examined the process of embryo and endosperm development in the mutant compared to WT. We found that the timing of embryo development differs slightly, with *crwn1/2* seeds exhibiting more embryos at earlier developmental stages at 5 days after pollination (DAP) and more variation in developmental stage at 7 DAP, ranging from globular embryos to bent cotyledon (Extended Data Fig. 2a). The variable embryo stages observed at 7 DAP might reflect the fact that *crwn1/2* seeds have mixed fates at maturity – seed abortion, small abnormal seeds, or wild-type sized seeds with mild abnormalities (Figs. 1a-d and 2a).

After fertilization the endosperm is initially coenocytic. Endosperm cellularization occurs around the heart stage of embryogenesis and is a key developmental transition point associated with the shift from endosperm proliferation to differentiation and maturation. The chalazal cyst, which is thought to be important for nutrient transfer from maternal tissue^26^, lies at the chalazal pole of the endosperm, does not undergo cellularization, and contains endoreduplicated nuclei. While observing embryos in *crwn1/2* seeds, we noticed unusually large endosperm chalazal cysts in some seeds (Fig. 3a and Extended Data Fig. 2b). We measured the area of the chalazal cyst and found that *crwn1/2* seeds have statistically larger chalazal cysts at 6 and 7 DAP, with greater variability in size than WT (Fig. 3b). We also checked if *crwn1/2* endosperm cellularization is perturbed. In wild-type seeds having late heart or early torpedo embryos, micropylar endosperm and peripheral endosperm are clearly cellularized (Fig. 3a). In contrast, *crwn1/2* seeds endosperm showed mixed states of cellularization at the same embryo stages ranging from no cellularization to full cellularization. Together these results demonstrate how CRWNs influence morphology in reproductive tissue and timely development of embryo and endosperm.

**Fig. 3.**
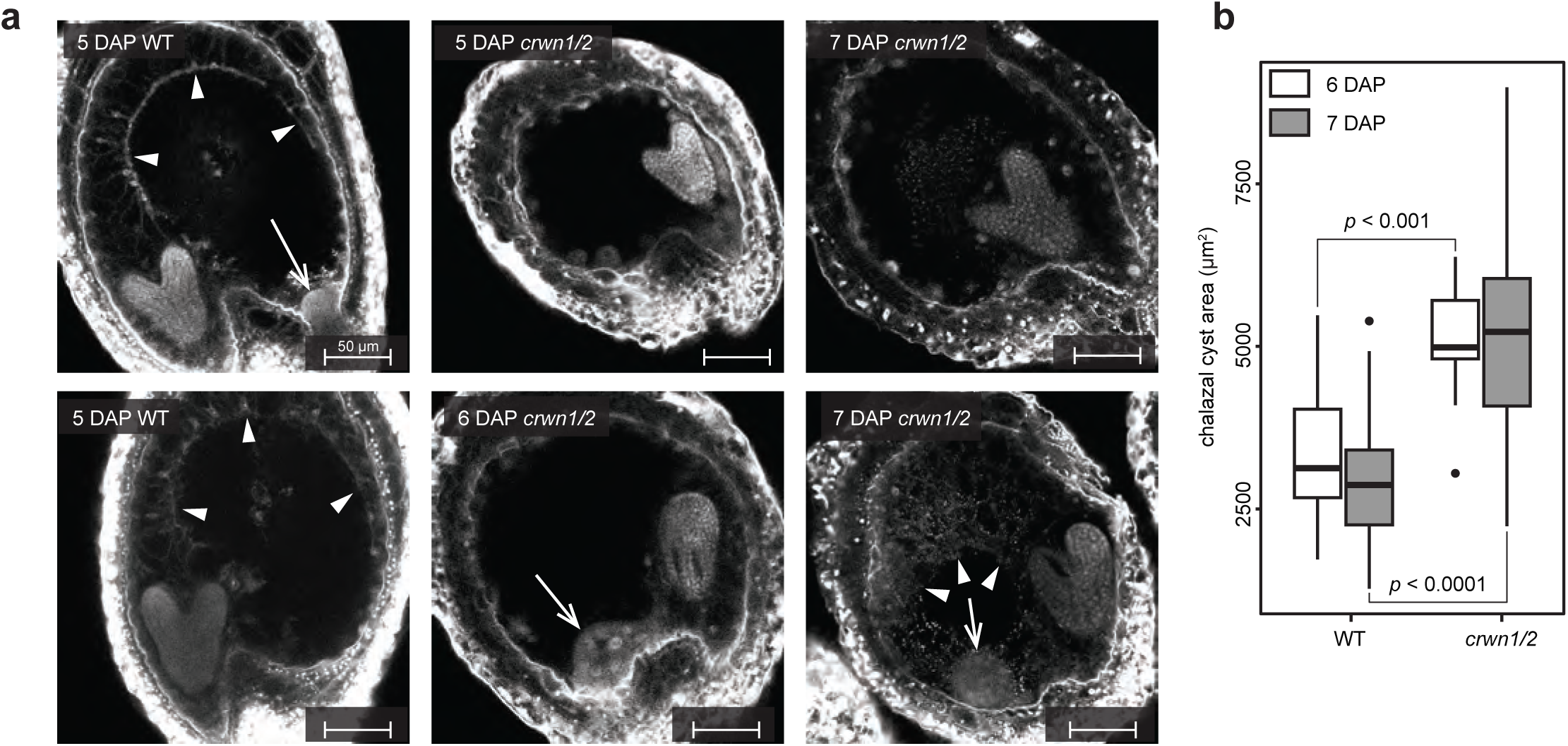
Enlarged endosperm chalazal cysts and endosperm cellularization delay in *crwn1/2* seeds. **a,** Confocal microscopy images of Feulgen-stained WT (left) and *crwn1/2* seeds at indicated days after pollination (DAP). Embryo stages were matched to enable comparison between WT and *crwn1/2*. Arrow heads, cellularized endosperm. Arrows, chalazal cysts. Scale bars, 50 μm. **b,** Comparison of chalazal cyst area between WT and *crwn1/2* seeds. Statistical analyses are Student’s *t*-tests between 6 DAP WT (*n* = 19) and 6 DAP *crwn1/2* (*n* = 11) and between 7 DAP WT (*n* = 40) and 7 DAP *crwn1/2* (*n* = 30). Boxplots show median (centerline), upper and lower quartile (Q3 and Q1. box), Q1 - 1.5*interquartile range (IQR) and Q3 + 1.5*IQR (whiskers), and outliers (points).

### Transcriptomic analyses reveal developmental mis-regulation and ectopic defense responses in *crwn1/2* mutant endosperm

After observing abnormal structural and developmental features in the endosperm, we characterized transcriptional differences between WT and *crwn1/2* endosperm to determine the molecular basis for these phenotypes. We performed mRNA-seq on *crwn1/2* and WT 6C endosperm nuclei that were isolated by FANS (fluorescence activated nuclei sorting) from whole seeds as previously described^27^ (Extended Data Fig. 3a-e). (The endosperm nuclei are triploid, whereas other seed nuclei are diploid.) We generated WT mRNA-seq profiles at 6 and 7 DAP and *crwn1/2* profiles at 6, 7, and 8 DAP, with 3 to 5 biological replicates per each genotype per time point. The gene expression profiles were highly correlated among biological replicates (Fig. 4a,b). Sample clustering was consistent with the slight developmental delay observed in *crwn1/2* seeds (Figs. 3a, 4a,b and Extended Data Fig. 2a). To minimize the identification of gene expression differences between WT and *crwn1/2* that were due to developmental mismatches, we focused on four gene expression comparisons: 6 DAP *crwn1/2* vs. 6 DAP WT (6vs6), 7 DAP *crwn1/2* vs. 6 DAP WT (7vs6), 7 DAP *crwn1/2* vs. 7 DAP WT (7vs7), and 8 DAP *crwn1/2* vs. 7 DAP WT (8vs7) (Fig. 4c, Extended Data Fig. 4a, and Supplementary Data 1).

**Fig. 4.**
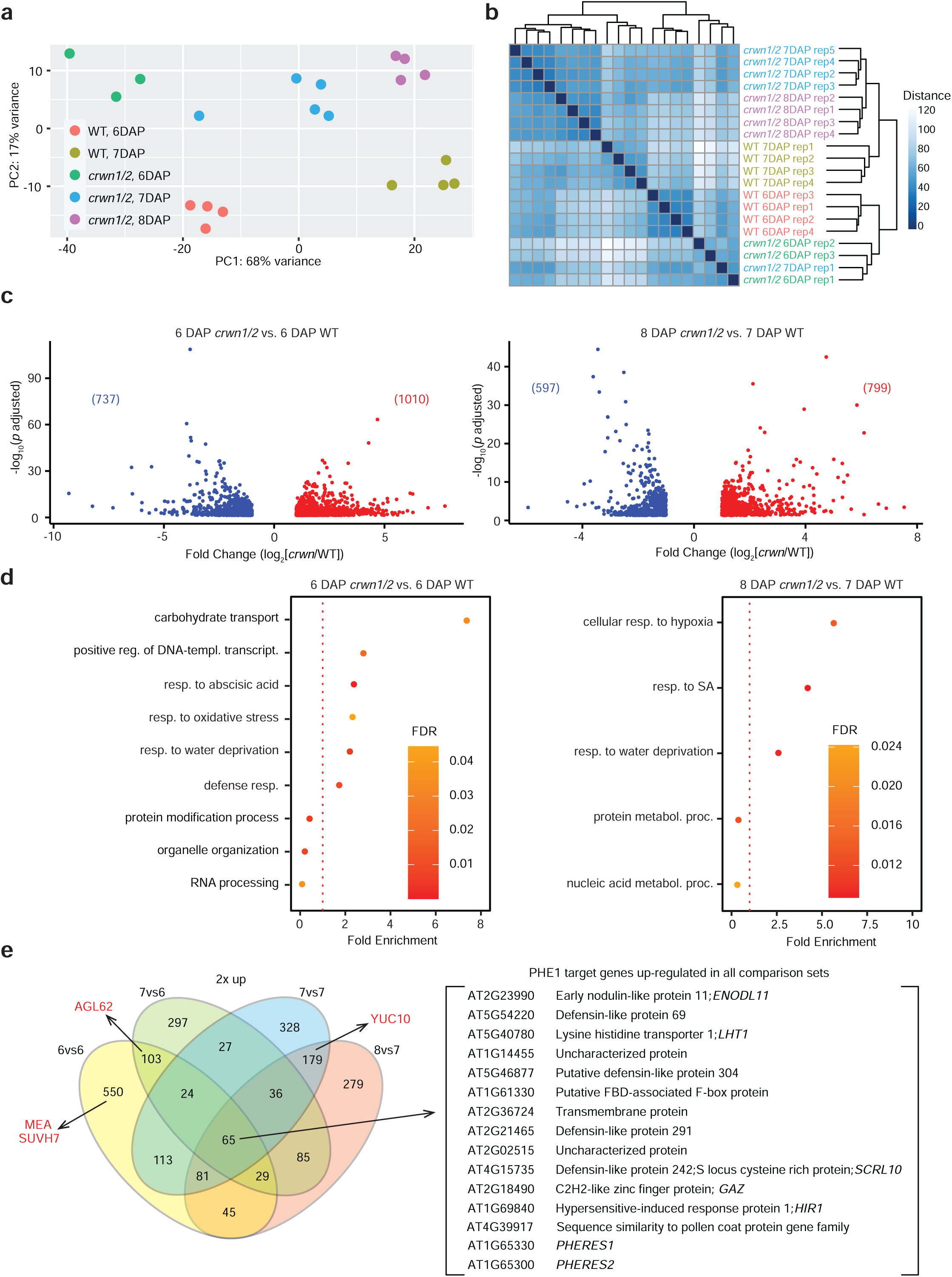
Transcriptome of *crwn1/2* endosperm reveals ectopic stress respons-es and increased expression of *PHE1* target genes. **a,** PCA plot of endosperm nuclear mRNA-seq samples. **b,** Heat map of the sample-to-sample distances among endosperm nulcear mRNA-seq samples. **c,** Volcano plots of endosperm nuclei mRNA-seq data comparing 6 DAP *crwn1/2* to 6 DAP WT (left) and 8 DAP *crwn1/2* to 7 DAP WT (right). Genes differentially expressed in *crwn1/2* (log_2_ >= 1 (red dots) or log_2_ <= -1 (blue dots), *padj* < 0.05) were plotted. **d,** GO overrepresen-tation test (PANTHER ’GO biological process complete’ set) of differentially expressed genes with increased expression in *crwn1/2* (log_2_ >= 2, *padj* < 0.05). The most specific sub-class GO terms are plotted. Dashed lines represent no (one fold) enrichment. Complete tables of enriched GO terms are available in Supple-mentary Data 1. **e,** Venn diagram of differentially expressed genes with increased expression in *crwn1/2* endosperm nuclei (log_2_ >= 1, *padj* < 0.05). 65 genes with increased expression in all comparison sets are enriched for 13 PHE1 target genes (3.24 fold enrichment in hypergeometric test, *p* < 0.001. See method.). Red-colored genes indicate PHE1 targets important in endosperm development as previously described by Batista et al. (2019). PHE1 target genes are described by the same authors.

We determined if differentially expressed genes in *crwn1/2* endosperm were enriched in any biological process. We found that stress response-related GO categories were shared among genes that were at least four-fold more highly expressed in *crwn1/2* endosperm (log2 >= 2, *p adjusted* < 0.05). These include cellular response to hypoxia, response to water deprivation, and defense or SA response-related GO categories (Fig. 4d and Extended Data Fig. 4b). As controls, we also compared 7 DAP WT to 6 DAP WT, 7 DAP *crwn1/2* to 6 DAP *crwn1/2*, and 8 DAP *crwn1/2* to 7 DAP *crwn1/2*. We checked GO overrepresentation among genes that were at least four-fold more highly expressed at the older stage (Extended Data Fig. 4c). In all comparisons, lipid storage and seed maturation GO terms (or a sub-class of seed maturation such as seed dormancy process) were found, which represents the normal course of seed development. Unlike inter-genotype comparisons, we did not identify defense response-related GOs. Thus, the differences in defense gene expression between WT and *crwn1/2* mutants do not simply reflect the normal developmental course of seed maturation but are due to disruption of *CRWN* genes. GO overrepresentation tests of genes that were down-regulated at least four-fold in *crwn1/2* mutants (log2 <= -2, *p adjusted* < 0.05) compared to WT showed no common GO terms among them (Supplementary Data 1)

We focused on genes whose expression increased by at least 2-fold in all four comparisons (log2 >= 1, *p adjusted* < 0.05) and identified 65 genes (Fig. 4e and Supplementary Data 1). Consistent with the GO overrepresentation tests, we found multiple defensin-like genes^28,29^ and stress-related genes (Supplementary Data 1) among the 65 genes. Additionally, we found that the Type I MADS box transcription factors *PHE1*/*AGL37* (*PHERES1*/*AGAMOUS-LIKE 37*) and *PHE2/AGL38* (*PHERES2*/*AGAMOUS-LIKE 38*) were highly up-regulated among the 65 genes, at least eightfold compared to WT endosperm (Fig. 4e and Supplementary Data 1). *PHE1* and *PHE2* are broadly expressed throughout WT endosperm prior to cellularization, with expression becoming more restricted to the chalazal endosperm after cellularization^30^. PHE1 endosperm targets have previously been defined by ChIP-Seq ^31^. We found that 13 genes commonly up-regulated in all four comparisons are known PHE1 targets, representing a 3.24-fold enrichment (hypergeometric test resulting *p* < 0.001. See method.)^31^. Among five defensin-like genes commonly up-regulated, four are PHE1 targets (Fig.4e and Supplementary Data 1). Additionally, PHE1 target genes involved in endosperm development^31^ were identified in some individual comparisons (Fig. 4e red), including *AGL62*, which suppresses endosperm cellularization^32^. In contrast, genes down-regulated in all four comparisons have only one PHE1 target (*DMT2*/*MET2*) (Supplementary Data 1). In addition, *PHE1* and *PHE2* are not expressed in *crwn1/2* leaves^19^, indicating that these genes are specifically up-regulated in the *crwn1/2* endosperm (Supplementary Data 1).

### H3K27me3 landscape is altered in *crwn1/2* endosperm and leaves

*PHE1* and *PHE2* expression is known to be repressed by the FIS2-PRC2 complex, which methylates histone H3 at lysine27, in endosperm^30,33,34^. The mRNA-seq results thus suggested that *crwn1/*2 mutations may disrupt endosperm H3K27me3 patterning. Other published data also suggest this possibility – the PRC2 interactor PWO1 (PWWP-DOMAIN INTERACTOR OF POLYCOMBS1) interacts with CRWN1 and regulates nuclear morphology^15,16^. Additionally, it was previously shown by H3K27me3 ChIP-qPCR that the defense response gene *PR1* (*PATHOGENESIS-RELATED GENE 1*), the SA-biosynthesis regulator *SARD1* (*SAR DEFICIENT 1*), and *CBP60g* (*CALMODULING-BINDING PROTEIN 60-LIKE*) lose H3K27me3 and increase in expression in *crwn1/2* leaves^19,20^. However, it is unknown if there is a genome-wide relationship between CRWNs and H3K27me3 in leaves or other tissues. Thus, we hypothesized that the *crwn1/2* endosperm phenotypes were caused at least in part by disruptions to H3K27me3 patterning in endosperm. Notably, *crwn1/2* mutants lack the completely penetrant seed abortion phenotype of FIS2-PRC2 mutants, suggesting that any alterations to H3K27me3 in *crwn1/2* are unlikely to be complete losses. To test whether *crwn1/2* plants have altered chromatin patterning, we produced H3K27me3 profiles of both endosperm and leaves in WT and *crwn1/2* using CUT&RUN.

We isolated 6C endosperm nuclei at 6 DAP for WT and 7 DAP for *crwn1/2* and performed CUT&RUN in biological triplicate with H3K27me3, H3 and IgG antibodies as previously described^27^ (Extended Data Fig. 3f,g). We chose two different time points to account for the developmental delay observed in *crwn1/2* (Figs. 3a, 4a, and Extended Data Fig. 2a). We performed the same profiling in WT and *crwn1/2* leaf nuclei to determine if any differences were specific to tissue type (Extended Data Fig. 3h,i). Pearson correlations of H3K27me3/H3 signals between samples showed that profiles were most similar by tissue type and then by genotype, suggesting that *crwn1/2* mutants retain an overall similar H3K27me3 landscape to WT (Extended Data Fig. 5a).

We identified 4,000 to 5,000 H3K27me3 peaks in each sample, with an average peak length of 3.5 and 2.6 kb in WT and *crwn1/2* endosperm and 4.8 and 4.6 kb in WT and *crwn1/2* leaves, respectively (Supplementary Data 2). Consensus peaks were defined as regions called as peaks in all three replicates in each tissue per genotype (Supplementary Data 2). We observed that endosperm H3K27me3 signal has a smaller dynamic range than in leaves, regardless of genotype (Fig. 5a). This may reflect the mixed epigenetic state present in endosperm, where it is known that the paternally-inherited genome has less H3K27me3 than the maternally-inherited genome, leading to overall hypomethylation ^35,36^. Although *crwn1/2* had a similar number of H3K27me3 peaks as WT, the average H3K27me3 peak height was lower in *crwn1/2* in both the endosperm and leaves (Fig. 5a). This suggests that there is a global dampening of H3K27me3 signal in *crwn1/2*. This was also observed when the average ratio of H3K27me3/H3 was plotted along the entire chromosome (Extended Data Fig. 6a,f).

**Fig. 5.**
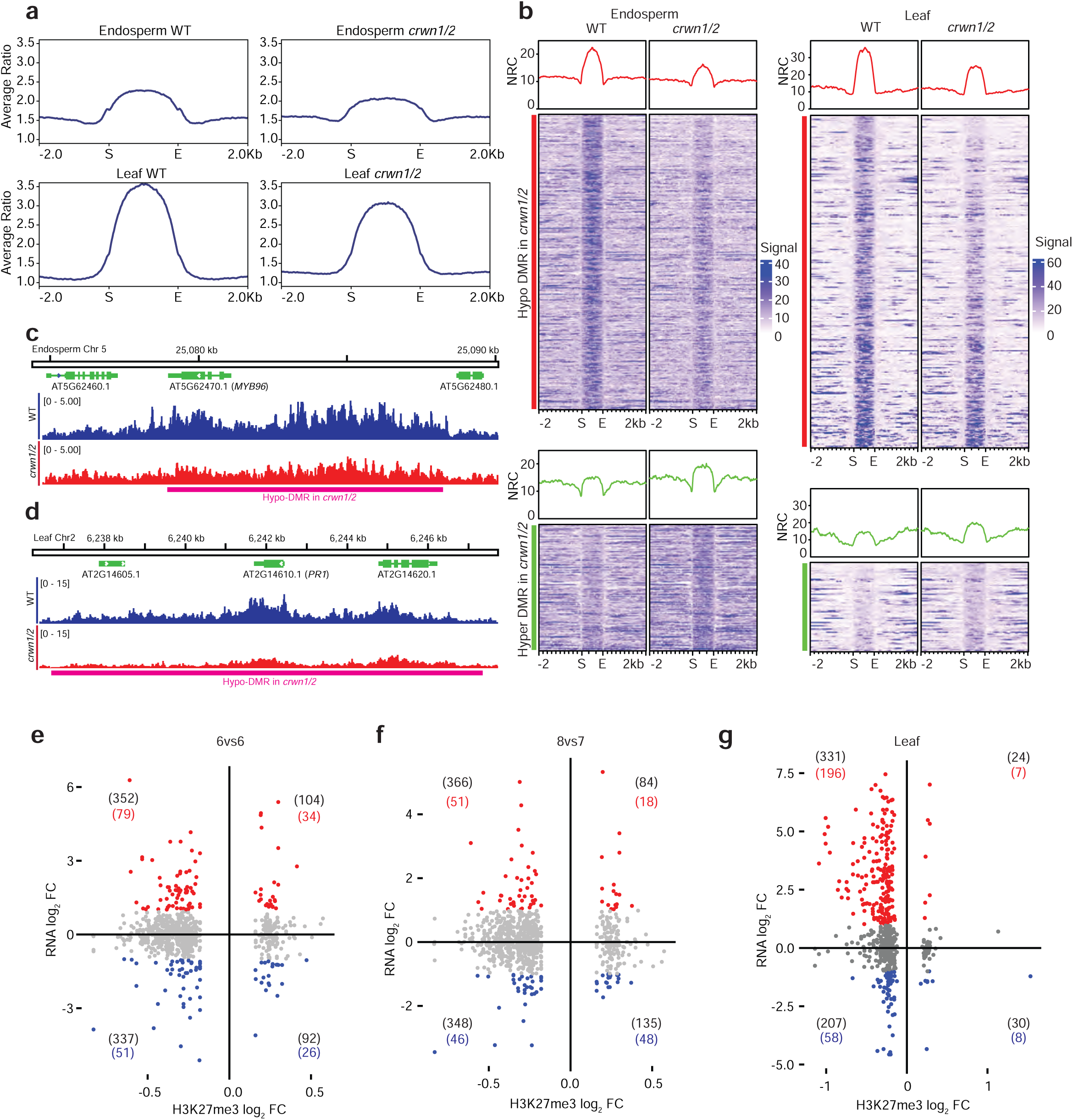
Both *crwn1/2* endosperm and leaves exhibit reduced H3K27me3 signals. **a,** Average H3K27me3/H3 ratio on H3K27me3 peak regions. Peak regions plotted in each genotype per tissue are consensus peaks among three biological replicates in each case. Individual peak loci are listed in Supplementary Data 2. **b,** H3K27me3 signals as DiffBind normalized read counts (NRC) of differentially H3K27-tri-methylated regions (DMRs) in *crwn1/2* mutants. Average H3K27me3 signals on hypo-(red line) and hyper-(green lines) DMRs are plotted at the top and heat map of individual DMRs are underneath. **c, d,** IGV browser snapshot of H3K27me3/H3 signal ratio on endosperm *MYB96* (**c**) and leaf *PR1* (**d**). Each track represents average H3K27me3/H3 signal ratio of three CUT&RUN replicates. Magenta boxes indicate DMRs. **e-g,** The differential expression of DMR genes as a log_2_ fold change (FC) in *crwn1/2* (regardless of *adjusted p* value) in the 6vs6 endosperm (**e**), 8vs7 endosperm (**f**), and leaf (**g**) RNA-seq comparisons is plotted against their log_2_FC in H3K27me3 levels. Numbers in parentheses are number of DMR genes in each quadrant (black, regardless of *adjusted p* value). RNA log_2_ FC >= 1 (red, regardless of *adjusted p* value), RNA log_2_ FC <= -1 (blue, regardless of *adjusted p* value).

We next asked if differentially H3K27-tri-methylated regions (DMRs) could be identified as distinct from the overall background of lower H3K27me3 in *crwn1/2*. Using DiffBind^37,38^, we identified 369 hypo-DMRs and 158 hyper-DMRs in *crwn1/2* endosperm (*FDR* <= 0.05) (Fig. 5b,c and Supplementary Data 3). These loci overlap with 1514 genes (endosperm DMR genes, defined by DMRs overlapping any portion of a gene), 1051 of which are hypomethylated and 463 are hypermethylated in *crwn1/2* mutants (Supplementary Data 3). Although *PHE1* and *PHE2* did not overlap a significant H3K27me3 DMR (Extended Data Fig. 5b), visual inspection of the region suggests lower H3K27me3 there. In leaves, 309 hypo-DMRs and 83 hyper-DMRs were detected (Fig. 5b,d and Supplementary Data 3). These loci overlap with 997 genes (leaf DMR genes), where 851 are hypomethylated and 146 are hypermethylated (Supplementary Data 3). Thus, in both tissues, the majority of the DMRs lose H3K27me3 in *crwn1/2* mutants, consistent with the genome-wide trend. DMR genes in endosperm and those in leaves are mostly unique to each tissue, with 135 DMR genes in common, (Extended Data Fig. 5c). This suggests that there are tissue-specific H3K27me3 level changes in *crwn1/2* mutants.

Hypo-DMR genes in *crwn1/2* endosperm are overrepresented for transcription-related GOs along with cutin and wax biosynthetic processes (Fig. 5c and Supplementary Data 3). In leaves, hypo-DMR genes are overrepresented with GO terms of defense and stress responses (Supplementary Data 3). These hypo-DMR genes include *WRKY55*^39^, and *CA10*^40^, which are involved in biotic and abiotic stress responses, respectively, as well as *PR1* (Fig. 5d and Extended Data Fig. 5b). Although *SARD1*^20^ was not identified as directly overlapping a hypo-DMR, a region 2 kb upstream (5’) was hypomethylated (Extended Data Fig. 5c).

Because H3K27me3 is known for its repressive roles in developmentally-responsive gene expression, we investigated if gene mis-regulation in *crwn1/2* correlated with DMRs. In the endosperm, changes in H3K27me3 level did not show a global correlation to changes in gene expression as less than 8% of DMR genes were differentially expressed (|log2| >= 1, *p adjusted* < 0.05) (Supplementary Data 4). In addition, hypo-DMR genes were associated with both increased and decreased expression in *crwn1/2* endosperm (Fig. 5e,f, Extended Data Fig. 5d,e, and Supplementary Data 4). In contrast, 22% of DMR genes in *crwn1/2* leaves were differentially expressed (|log2| >= 1, *p adjusted* < 0.05) (Supplementary Data 4). In addition, hypo-DMRs in *crwn1/2* leaves preferentially overlapped with genes with increased expression in *crwn1/2* leaves (Fig. 5g and Supplementary Data 4). Among these, the previously mentioned *PR1*, *SARD1*, *CA10* and *WRKY55* lost methylation and increased in expression (Supplementary Data 4), supporting the idea that ectopic defense responses in *crwn1/2* leaves correlate with the decrease of H3K27me3 level at those loci.

Together, these data indicate that CRWNs are involved in maintenance of tissue-specific H3K27me3 profiles. We also found that DMRs in the *crwn1/2* endosperm are enriched in transcription factor loci, implying that CRWNs might be involved in regulation of transcription factor genes in that tissue. In leaves, DMRs in *crwn1/2* mutants are enriched for defense-related genes, supporting the idea that CRWN1 and CRWN2 are necessary to maintain epigenetic control of defense gene regulation.

## Discussion

We investigated the involvement of CRWNs in gametophyte and seed development, which has not been extensively documented previously. After observing morphological and developmental defects in the endosperm of *crwn1/2* mutants, we also aimed to elucidate the underlying mechanisms of how CRWNs function in endosperm. We compared these results to similar experiments in leaves to elucidate any distinct impacts of *CRWNs* in different tissues.

Our initial observations revealed that *crwn* seeds display morphological and developmental defects. Altered development was most clear in endosperm chalazal cysts of *crwn1/2* seeds, which was accompanied by mild variations in the timing of embryo development and endosperm cellularization. Considering that the gametophyte generation precedes fertilization, we reasoned that abnormalities in *crwn* gametophtyes or the tissues bearing *crwn* gametophytes could potentially explain some of the post-fertilization seed phenotypes or reduced seed set in *crwn1/2* mutants. We found that *crwn* mutants have reduced ovule numbers and aberrant pollen morphologies. In addition, crosses revealed that the combined *crwn1/2/3/4* mutations were inherited through the male or female gametophyte at a significantly lower frequency than expected. Collectively, these findings support the essential roles of CRWNs in successful reproduction and the development of a full set of viable seeds.

We hypothesized that the developmental defects observed in *crwn* mutant seeds might be accompanied by endosperm-specific gene mis-regulation and epigenetic changes, which could be distinct from effects observed in vegetative tissues. CRWNs are required for nuclear differentiation and overall chromatin structure^11,41^. However, the endosperm has a distinct chromatin structure compared to leaves or seedlings, with relatively decondensed chromocenters ^42^ and parent-of-origin specific histone modifications^35^. Thus, the endosperm might display a distinct sensitivity to *CRWN* disruption. Our endosperm RNA-seq analysis revealed hundreds of differentially expressed genes in the *crwn1/2* mutant 6C seed nuclei. 6C nuclei are expected to be enriched for endosperm nuclei in S-phase and endosperm nuclei undergoing endoreplication, a feature primarily associated with chalazal endosperm^43^. The RNA-seq data uncovered two notable features of CRWN1’s and CRWN2’s roles in the endosperm. First, a considerable number of stress-related genes associated with both biotic and abiotic responses are mis-regulated in *crwn1/2* endosperm. This is consistent with the well-documented roles of CRWN1 and CRWN2 in defense responses in leaves and seedlings^18–21^. Previous studies demonstrated that *crwn1/3* mutants exhibit hypersensitivity to abscisic acid during seed germination, which is attributed to the role of CRWN3 in the degradation of ABI5^44^. However, it remained unknown whether similar phenomena occur in *crwn1/2* mutant seeds. The transcriptomic data of *crwn1/2* endosperm indicates that abiotic stress responses are also mis-regulated in the mutant endosperm, in addition to biotic stress responses. GO terms related to hypoxia and oxidative stress responses are also overrepresented, which suggests that reactive oxygen species (ROS) ^45–47^ might drive changes in in *crwn1/2* mutants. Additionally, the MADS-box transcription factor *PHE1* and its targets are mis-regulated in *crwn1/2* endosperm. *PHE1* is also highly expressed in the endosperm of paternal excess crosses, when diploid females are pollinated by tetraploid males. This is accompanied by delayed endosperm cellularization and enlarged chalazal cysts^26,48,49^, with some similarity to that observed in *crwn1/2* seeds.

Stress responses and developmental signals are likely interconnected in endosperm. For example, PHE1 target genes with increased expression in *crwn1/2* endosperm include *HIR1*^50,51^, defensins, and the GA- /ABA-responsive zinc finger protein (*GAZ*; AT2G18490)^52,53^, which are involved in immune responses (*HIR* and defensins) and hormone-related seed germination (*GAZ*). In addition, differentially expressed ROS-related genes in *crwn1/2* endosperm suggests that defense signaling and developmental signaling could be linked through CRWNs. The endosperm is subject to programmed cell death as a normal part of its development. ROS-involved programmed cell death, which is important for both tissue development and hyper-sensitive responses, could be related to CRWN’s functions in endosperm development.

How is this gene mis-regulation triggered? One compelling culprit is disruption of the H3K27me3 mark, based on three lines of prior evidence: by ChIP-qPCR ^19,20^ individual genes misregulated in *crwn1/2* leaves lose H3K27me3; CRWN1 interacts with the PRC2 interacting protein PWO1^15,16^; and *PHE1* expression is repressed by the PRC2 components MEA and FIE. We found that the level of H3K27me3 is reduced in both endosperm and leaves in *crwn1/2* mutants, consistent with and extending previous findings. We also found that hypo-DMR genes tend to be up-regulated in *crwn1/2* leaves, supporting the idea that the lamina contributes to a repressive gene expression environment. The correlation was not present, however, in *crwn1/2* endosperm. Because hypo-DMRs in *crwn1/2* endosperm coincide with many transcription factor genes (Supplementary Data 3), it is possible that the *crwn1/2* transcriptomics are more reflective of downstream processes resulting from the action of those transcription factors than what is observed in *crwn1/2* leaves. Alternatively, the lack of a strong connection between transcriptional changes and H3K27me3 in *crwn1/2* endosperm could reflect the variable penetrance of the *crwn1/2* phenotype, with some seeds ultimately aborting and some appearing mostly normal. Our profiles of developing seed endosperm at 7 days after pollination will reflect a mixture of seed phenotypes. It is also possible that the tissue-specific cellular environment affects gene expression patterns in *crwn1/2* endosperm in a manner distinct from that of leaves. For example, it is unknown if protein interactors of CRWNs described in vegetative tissues also interact with CRWNs in endosperm. Our data suggest that there are tissue-specific effects of CRWNs in development and epigenetic gene regulation. This may be more pronounced in endosperm because of its unusual chromatin landscape compared to other tissues.

The tissue-specificity of the lamina is well described in animals, where LADs consist of facultative and constitutive domains^54–60^. While constitutive LADs are invariable among cell types, the facultative LADs are defined in a cell-type specific manner. However, it is unknown if the plant lamina has an equivalent of facultative LADs. We found that plant lamina associated domains, which have been defined in seedlings as regions where CRWN1 interacts with the DNA^17^, could be divided into clusters based on their H3K27me3 levels in leaves (Extended Data Fig. 6a-d). In this *k*-means clustering (*k* = 3), the cluster with higher H3K27me3 levels (cluster 1) has the fewest number of differentially expressed genes in *crwn1/*2 leaves compared to the other clusters (Extended Data Fig. 6d,e). In addition, the numbers of down-regulated genes and up-regulated genes are similar in cluster 1 compared to clusters 2 and 3 (Extended Data Fig. 6e). Clusters with lower H3K27me3 levels (clusters 2 and 3) have more up-regulated genes than down-regulated genes in *crwn1/2* leaves (Extended Data Fig. 6d,e). We interpret this result as follows: PLAD genes in WT leaves are silenced by both the lamina-DNA interactions and H3K27me3 in cluster1, whereas PLAD genes in cluster 2 and cluster 3 rely more on the lamina-DNA interactions for gene silencing than on H3K27me3 (which is already low in WT). Thus, when the CRWN1-PLAD interactions are absent or disrupted in *crwn1/2* mutants, expression of PLAD genes is dependent on the level of H3K27me3: cluster 1 in *crwn1/2* leaves lacks the CRWN1-PLAD interaction but retains relatively high levels of H3K27me3, hence the low number and ratio of genes up-regulated, whereas cluster 2 and cluster 3 lack both the lamina-DNA interaction and H3K27me3, hence the more genes up-regulated in these clusters (Extended Data Fig. 6d,e). We view this phenomenon as a ’fail-safe’ to ensure gene silencing, so that the silenced genes are only expressed when neither H3K27me3 nor lamina-DNA interactions are present. When we apply vegetative-tissue-defined PLADs to the endosperm, however, genes in PLADs were mis-regulated equally in both directions, regardless of H3K27me3 levels in each cluster (Extended Data Fig. 6f-j). This suggests that PLADs defined in one tissue might not be readily applicable to another tissue in Arabidopsis.

The importance of this study lies in the demonstration that plant lamin analogs, CRWNs, have critical roles in efficient reproduction and in maintaining tissue specific epigenetic patterns and gene expression programs. We provided evidence in support of the long-suggested idea that CRWNs contribute to genome-wide H3K27me3 levels and patterns, which is consistent with the idea that the nuclear periphery is in general a repressive gene expression environment where the lamina contributes to gene silencing. The essential roles of CRWNs in Arabidopsis leave a clue as to why plants evolved the nuclear lamina. CRWN genes and lamin genes are not homologous, yet mounting knowledge of the plant lamina argues that eukaryotic nuclei tend to develop this reticular structure to better control their genomes and support nuclear structure. Investigating *crwn* mutants gives comparative biological understandings of the plant lamina, which highlights the sub-nuclear level of convergent evolution in eukaryotes.

## Methods

### Plant materials, accessions, and primers used for genotyping

All plants except ones for germination tests were grown at approximately 22°C in a greenhouse on soil under long day conditions (16 h light/8 h dark). Each of single *crwn* mutant lines used to make *crwn1/2/4* and *crwn1/3/4* were originally from the Arabidopsis Biological Resource Center at Ohio State University and are described by Wang et al^12^. T-DNA line accessions for these mutant alleles are SALK_025347 (*crwn1-1*), SALK_076653 (*crwn2-1*), SALK_099283 (*crwn3-1*), and SALK_079296 (*crwn4-1*). The *sid2-1* mutant allele is from the Metraux group^61^. Primers used to confirm genotypes of mutant alleles are listed in Supplementary Data 6.

### Germination test

WT and *crwn1/2* mutant plants were grown together under the same greenhouse conditions described above and seeds were harvested simultaneously. Harvested seeds were defined as small or large based on whether they passed through a mesh with a 0.2328μm opening. Seeds were sterilized with 75% ethanol for 30 sec and 50% bleach for 30 sec, rinsed, resuspended in distilled sterile water, and plated on 0.5x Murashige and Skoog (MS) agar plates with 1% sucrose. MS plates were stratified for 2 days in 4°C. Seeds were marked as germinated if the radicle was visible. Seed germination was counted every other day.

### Pollen nuclei migration pattern

Pollen grains were germinated as described in Boavida and McCormick (2007)^62^. Briefly, the 1.5% low melting agar with pollen germination solution was poured on microscope slides and pollen grains were put on the cooled surface. The slides were stored in a humid box until the pollen germinated. Germinated pollen grains were stained with Hochest 33342 solution described in Goto et al. (2020)^24^. Samples were observed using Zeiss Axio Imager M2.

### Scanning electron microscopy (SEM)

Seed and pollen samples for SEM were submitted to Peterson Nanotechnology Materials Core Facility at the Koch Institute’s Swanson Biotechnology Center. Briefly, pollen grains were collected by rubbing anthers on conductive adhesive carbon tape attached to SEM pin stubs (Ted Pella, Inc Product #16084-1 and #16111). Samples were then sputter coated with gold or platinum in a Denton Desk V coater and transferred to a Zeiss Crossbeam 540 field emission SEM. Observations were made at varying accelerating voltages.

### Feulgen staining and confocal microscopy

Siliques were slightly cut open along the dehiscence zone between replum and valves using a sharp needle in order to ensure penetration of solution into siliques. Open siliques were stained as described by Braselton et al.^63^ using LR white acrylic resins (Sigma-Aldrich #L9774) and Schiff’s reagent (Sigma-Aldrich #3952016). Samples were observed with Zeiss LSM 710 NLO Laser Scanning Confocal in W.M. Keck Microscopy Facility at Whitehead Institute.

### Seed light microscopy and chalazal cyst measurements

Seeds were cleared with 70% (w/v) chloral hydrate solution. Specimens were observed and staged using Zeiss Axio Imager 2 microscopy with a differential interference contrast (DIC) filter. When visible, chalazal cyst area was determined by measuring the length and width of each cyst using tools in the Zeiss ZEN software.

### Nuclei extraction and fluorescence-activated nuclei sorting

For endosperm CUT&RUN, 6 DAP WT seeds or 7 DAP *crwn1/2* seeds from 4 to 7 siliques were pulverized with a pestle (AXYGEN #PES-15-B-SI) in a 1.5mL tube containing 100μL CyStain UV Precise P Nuclei Extraction Buffer (Sysmex #05-5002-P02). 800μL staining buffer (Sysmex #05-5002-P01) was added and mixed. The nuclei solution was filtered with two CellTrics 30μm disposable filters (Sysmex #04-004-2326) in tandem. The filtered solution was sent to Whitehead Flow Cytometry Core Facility and sorted by BD FACSAria II using 100μm nozzle, 2500 plates voltage and 20 sheath pressure setups. This process was repeated several times to acquire enough 6C nuclei. For leaf CUT&RUN, 2 to 3 of 5th leaves were first chopped with a clean razor blade on a petri dish with 150μL of the nuclei extraction buffer and then 1200μL of the staining buffer was added after the chopping. The mixture was filtered in the same way as the endosperm. The flow cytometry sorting procedure is the same as the endosperm CUT&RUN. For endosperm mRNA-seq, seeds from 1 to 5 of siliques were used with the same methods as in the endosperm CUT&RUN. The numbers of siliques used and nuclei collected for each sample are available in Supplementary Data 6.

### mRNA-seq library preparation, sequencing, and data analysis

6C seed nuclei sorted by FANS described above were used for RNA-seq. Library was prepared following the protocols provided by the manufacturer in SMART-Seq mRNA LP kit (Takara Bio #634768) and Unique Dual Index Kit (U001-U024) (Takara Bio #634756) with some modifications. Although the manufacturer recommended cDNA synthesis with up to 1000 cells and no indication for nuclei number, we used up to 3000 nuclei, which did not hinder the downstream procedure. The cDNA and library were purified as indicated in the manufacturer’s protocol with AMPure XP beads (Beckman Coulter #A63881). After the library amplification and purification, we checked the fragment length using an Agilent BioAnalyzer 2100. If there were leftover primer-dimers or other small fragments detected, additional purification with 1:1 bead to sample volume was performed and re-analyzed by the BioAnalyzer. The purified samples were sequenced by Whitehead Genome Technology Core using NovaSeq 6000 and NovaSeq SP v1.5 reagents. Paired end read lengths were 50 bp by 50 bp.

FASTQ files were trimmed with trim_galore (version 0.6.7) with ’-q 25’ and ’--stringency 3’ options. To align trimmed sequences, genome files for STAR (version 2.7.1a) were first generated using STAR ’genomeGenerate’ mode and TAIR10 whole genome sequence with ’--genomeSAindexNbases 12’ and ’-- sjdbOverhang 30’ options. Trimmed sequences were aligned to the genome files using STAR with ’-- alignSoftClipAtReferenceEnds No’, ’--alignEndsType EndToEnd’, ’--outSAMstrandField None’, ’-- runThreadN 1’, ’--outFilterType BySJout’, ’--outFilterMultimapNmax 5’, ’--outFilterMismatchNmax 10’, ’-- outFilterMismatchNoverReadLmax 0.05’, ’--alignIntronMin 70’, ’--alignIntronMax 5000’, ’-- alignMatesGapMax 100000’, and ’--outFilterIntronMotifs RemoveNoncanonical’ options. PCR duplicates were removed using Picard MarkDuplicates (version 1.121). These trimmed and de-duplicated reads were indexed by Samtools^64^ (version 1.11) and counted by htseq-count^65^ (version 1.99.2) with ’-- mode=intersection-nonempty’ and ’--nonunique=all’ options. Differential expression analysis was performed using DESeq2^66^ (version 1.36.0) with ’DESeqDataSetFromHTSeqCount’ function. We included genes only when the gene had at least 10 counts and at least three samples met this condition, using ’keep <- rowSums(counts(deseqdataset) >= 10) >= 3’. For leaf RNA-seq, this was changed to 6 counts and 2 samples as there were fewer replicates than for endosperm.

### PCA and clustering analysis

To generate PCA plots from endosperm nuclear mRNA-seq samples (Fig. 4a), DESeq2 variance stabilizing transformation (*vst*) function was applied to read counts generated by HTSeq (htseq-count) and the resulting *vst* object was used for plotting. For distance clustering (Fig. 4b), the *vst* object was used to calculate sample-to-sample distances using *dist* function. For Pearson correlation matrix of H3K27me3/H3 signals of all CUT&RUN samples (Extended Data Fig. 5a), we used ’plotCorrelation’ from deepTools.

### GO analysis

PANTHER (https://pantherdb.org/) ’statistical overrepresentation test’ was used to find GO terms prevalent in list of genes^67,68^. For test type and correction, default setting of Fisher’s exact test and calculating false discovery rate were used.

### Hypergeometric test

To test if PHE1 targets are statistically overrepresented among 65 up-regulated genes in all four RNA-seq comparisons (Fig. 4e), we employed hypergeometric test with population size 27,430 (PANTHER mappable IDs), number of successes in the population 1,696 (PHE1 targets), samples size 65 (up-regulated genes in all four RNA-seq comparisons), and number of successes 13 (PHE1 targets among 65 genes). The result gave over enrichment of 3.24 fold compared to expected number of successes of 4.01 with *p* < 0.001.

### CUT&RUN library preparation, sequencing, and data analysis

6C nuclei in seeds were sorted by FANS as above and used for CUT&RUN. CUT&RUN procedure and library preparation protocol used in this study is described by Zheng and Gehring 2019. We used IDT for Illumina - TruSeq DNA UD indexes. The library was checked with Agilent BioAnalyzer 2100 at Whitehead Genome Technology Core to ensure no primer dimers were remaining in the samples. The library was sequenced using NovaSeqSP 6000 and NovaSeq SP v1.5 reagents with 50 bp by 50 bp paired end setup in Whitehead Genome Technology Core.

CUT&RUN sequencing data were processed by Epic2 (version 0.0.52), DiffBind (version 3.8.4), deepTools^69^ (version 3.4.3) and KaryoploteR^70^ (version 1.24.0). FASTQ files were trimmed with trim_galore (version 0.6.7) with ’--stringency 3’, ’-r1 32’, ’-r2 32’, ’-q 20’, ’--clip_R1 5’, and ’--clip_R2 5’ options. Trimmed data were aligned to Arabidopsis genome by Bowtie2 (version 2.3.4.1) with ’-L22’, ’-N 0’, and ’-X 1000’ options. Uniquely mapped reads were extracted by Samtools with ’view -q3’ option and Picard MarkDuplicates was used to remove PCR duplicates. Epic2 peak caller^71^ was used to define H3K27me3 peaks. The reads were passed to Epic2 using H3 reads as control. ’--effective-genome-fraction’ was set as 1 and bin size was 150. To find DMR between WT and *crwn1/2* mutants, DiffBind was used with H3 reads as controls and IgG targets as greylists. Reads were counted using ’dba. count’ function with ’summits = TRUE’. DMR genes were defined as genes overlapping any portion of it with DMR. deepTools bamcompare was used to generate average H3K27me3/H3 ratio on genomic regions in Fig. 5a and Extended Data Fig. 6a,c,e,g with ’--operation ratio’, ’--scaleFactorsMethod readCount’, ’--skipNonCoveredRegions’, and ’-bs 20’ options.

### *k*-means clustering of PLADs based on H3K27me3

Bigwig files (H3K27me3/He) generated by deepTools in CUT&RUN data analysis were averaged by WiggleTools^72^ ’mean’ mode. deepTools ’computeMatrix’ files per tissue were generated by inputting both genotypes at one computeMatrix process. Resulting computeMatrix files of each tissue were used to generate cluster profiles by deepTools ’plotProfile’ mode with ’--kmeans’ 1 (no clustering) or 3 (3 clusters) and ’--perGroup’ options.

### Data availability

H3K27me3 CUT&RUN data (for both 6C endosperm nuclei and 2C-16C leaf nuclei) and mRNA-seq data (for 6C endosperm nuclei) are available at NCBI Sequence Read Archive (NCBI GSE 243032, BioProject ID PRJNA1006063, BioSample Accessions SAMN37008819 - SAMN37008874) and Supplementary Data. Leaf mRNA-seq data are available as NCBI BioProject ID PRJNA485018^19^.

## Acknowledgements

We would like to express our gratitude for scientific services and technical support provided by Sumeet Gupta and Amanda Chilaka at the Whitehead Genome Technology Core, Patrick Autissier and Aditya Rathee at the Whitehead Flow Cytometry Core, Cassandra Rogers and Asier Marcos Vidal at the Whitehead W.M. Keck Microscopy Facility, and Abigail Lytton-Jean and David Mankus at the Peterson (1957) Nanotechnology Materials Core Facility in the Koch Institute’s Robert A. Swanson (1969) Biotechnology Center. We are grateful to Eric J. Richards, Jay Goodman, and Roman Podolec for their valuable feedback on the manuscript. For CUT&RUN data analysis, we acknowledge help from Xiao-yu Zheng and Colette Picard. This research was supported by NIH R35GM145321.

---

**Extended Data Fig. 1.**
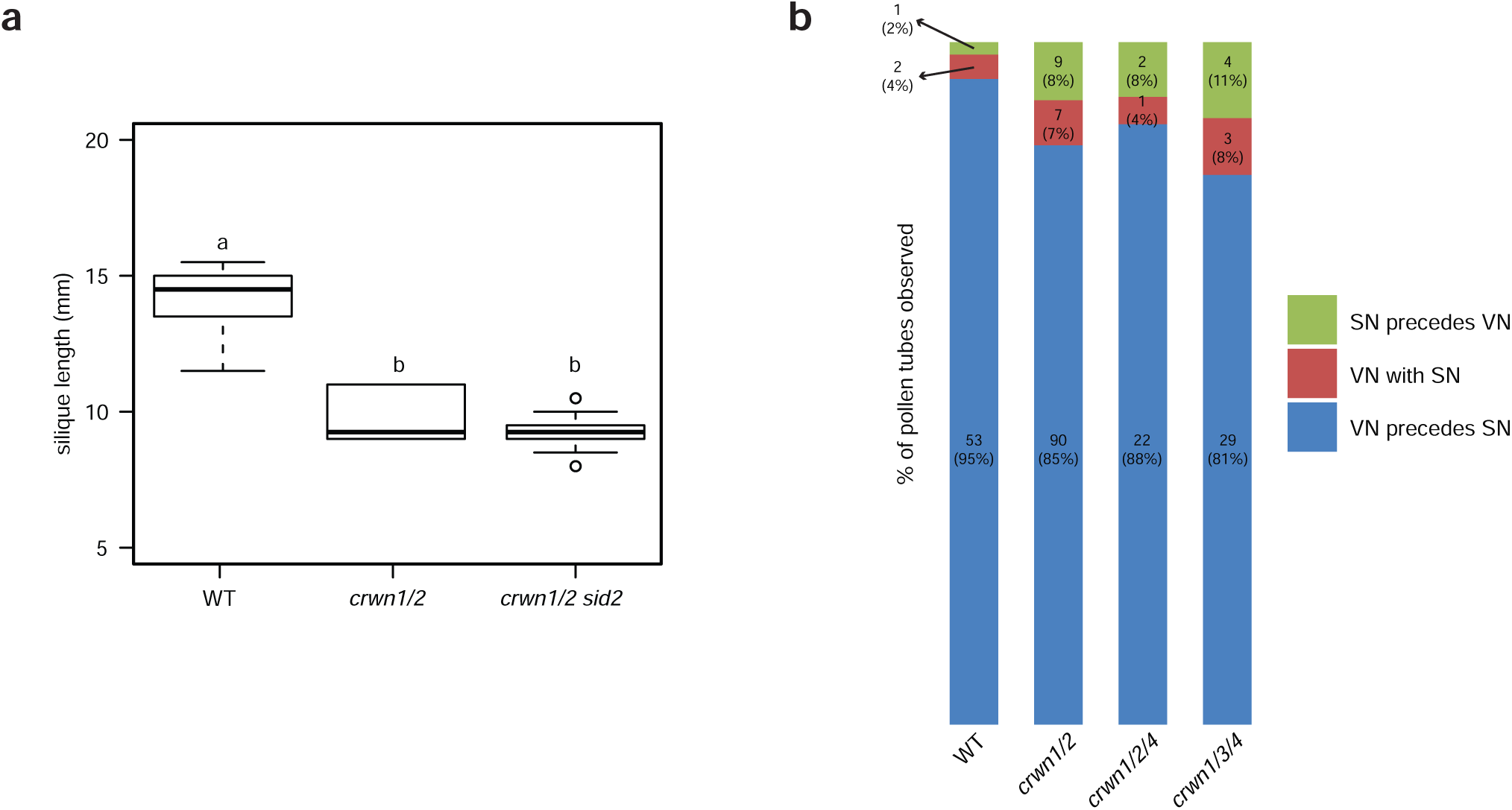
Length of *crwn1/2 sid2* mutant siliques and pollen nuclei migration pattern in *crwn* mutants. **a,** Length of 10 siliques at the yellowing stage was measured for each genotype of plants grown under the same conditions. Note that this experiment is independent from Fig. 1b. Boxplots show median (centerline), upper and lower quartile (Q3 and Q1. box), Q1 - 1.5*inter-quartile range (IQR) and Q3 + 1.5*IQR (whiskers), and outliers (points). Statistical analyses are one-way ANOVA with post hoc Tukey’s tests. Different letters indicate significant differences among them (*p* < 0.05). **b,** The pollen nuclei migration pattern in *crwn* mutants is similar to the WT. Pollen grains were germinated on an agar plate with pollen germination media in a humid environment (see Methods). SN, sperm cell nuclei. VN, vegetative cell nucleus.

**Extended Data Fig. 2.**
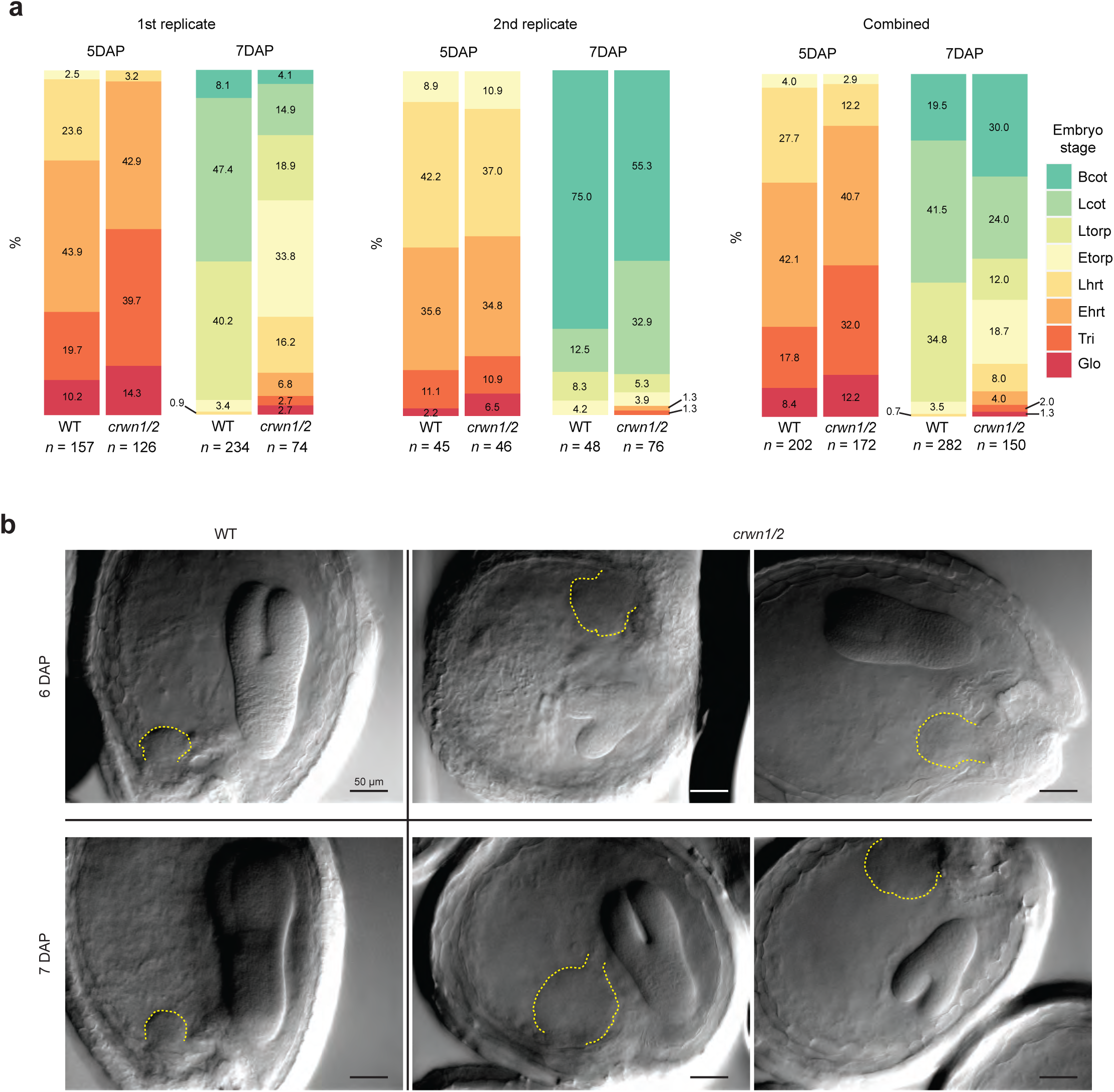
Embryo and endosperm development in *crwn1/2* mutants. **a,** Seeds at 5 DAP and 7 DAP were cleared in a chloral hydrate solution, and the embryo stages were visually determined. The 1st replicate (left) and 2nd replicate (center) were independently observed. A single plot from the two replicates combined is also shown (right). Glo, globular embryo stage. Tri, triangular. Ehrt, early heart. Lhrt, late heart. Etorp, early torpedo. Ltorp, late torpedo. Lcot, linear cotyledon. Bcot, bent cotyledon. **b,** Chalazal cysts (demarcated by dashed yellow lines) of WT and *crwn1/2* seeds at 6 DAP and 7 DAP. Seeds were cleared in a choloral hydrate solution and observed using Differential Interference Contrast (DIC). Scale bars, 50μm.

**Extended Data Fig. 3.**
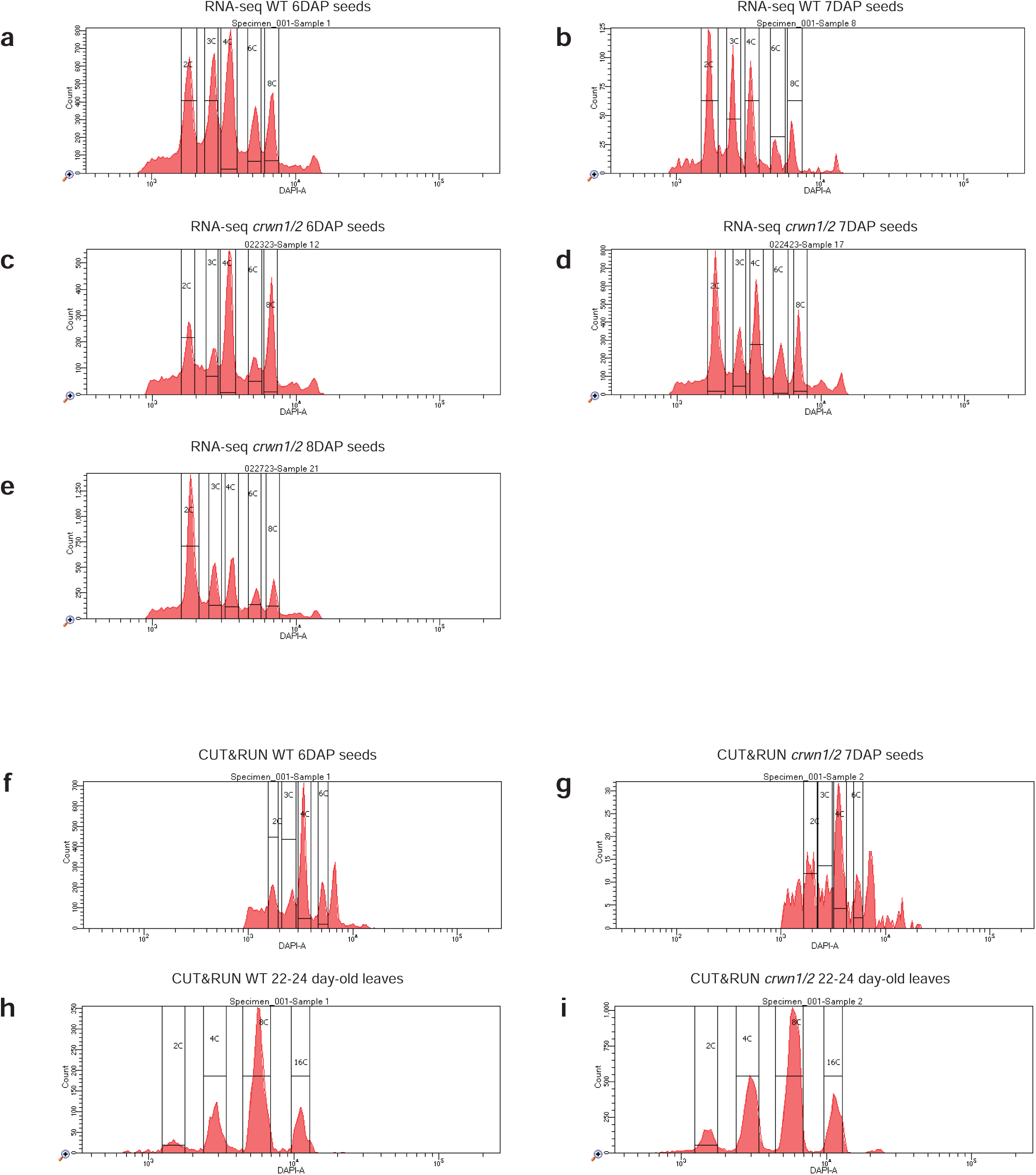
Representative images of FANS peaks for mRNA-seq and CUT&RUN. Nuclei were stained with DAPI solution and then sorted by their ploidy (see Methods). For endosperm mRNA-seq, 6C nuclei in 6DAP WT **(a)**, 7DAP WT **(b)**, 6DAP *crwn1/2* **(c)**, 7DAP *crwn1/2* **(d)**, and 8DAP *crwn1/2* **(e)** seeds were sorted. For endosperm CUT&RUN, 6C nuclei in 6DAP WT **(f)** and 7DAP *crwn1/2* **(g)** seeds were sorted. For leaf CUT&RUN, 2C - 16C nuclei in 5th leaves from 22-24 day-old WT **(h)** and *crwn1/2* **(i)** plants were sorted.

**Extended Data Fig. 4.**
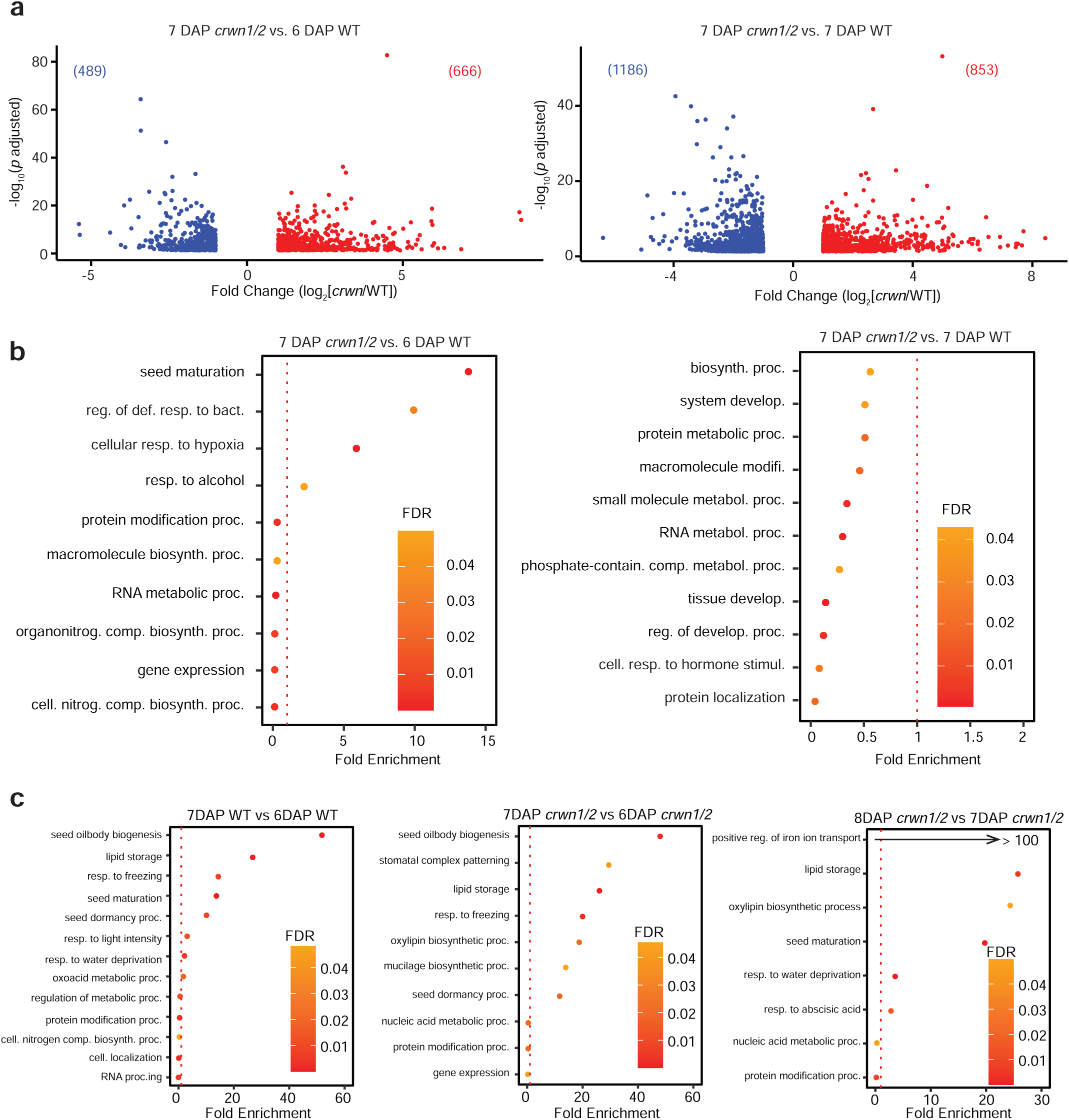
Differentially expressed genes in *crwn1/2* endosperm nuclei and GO overrepresentation tests. **a,** Volcano plots of differentially expressed genes (log_2_ >= 1 (red dots) or log_2_ <= -1 (blue dots), *p adjusted* < 0.05) in *crwn1/2* endosperm, comparing 7 DAP *crwn1/2* to 6 DAP WT (left) and 7 DAP *crwn1/2* to 7 DAP WT (right). Numbers in brackets indicate the number of genes in the same color category. **b, c,** GO overrepresentation test (PANTHER ’GO biological process complete’ set) of differentially expressed genes with increased expression in *crwn1/2* endosperm nuclei (log_2_ >= 2, *p adjusted* < 0.05) compared to WT (**b**) and differentially expressed genes with increased expression in older stages (log_2_ >= 2, *p adjusted* < 0.05) compared to younger stages of the same genotype (**c**). The most specific sub-class GO terms are plotted. Dashed lines represent no (one fold) enrichment. Complete tables of enriched GO terms are available in Supplementary Data 1.

**Extended Data Fig. 5.**
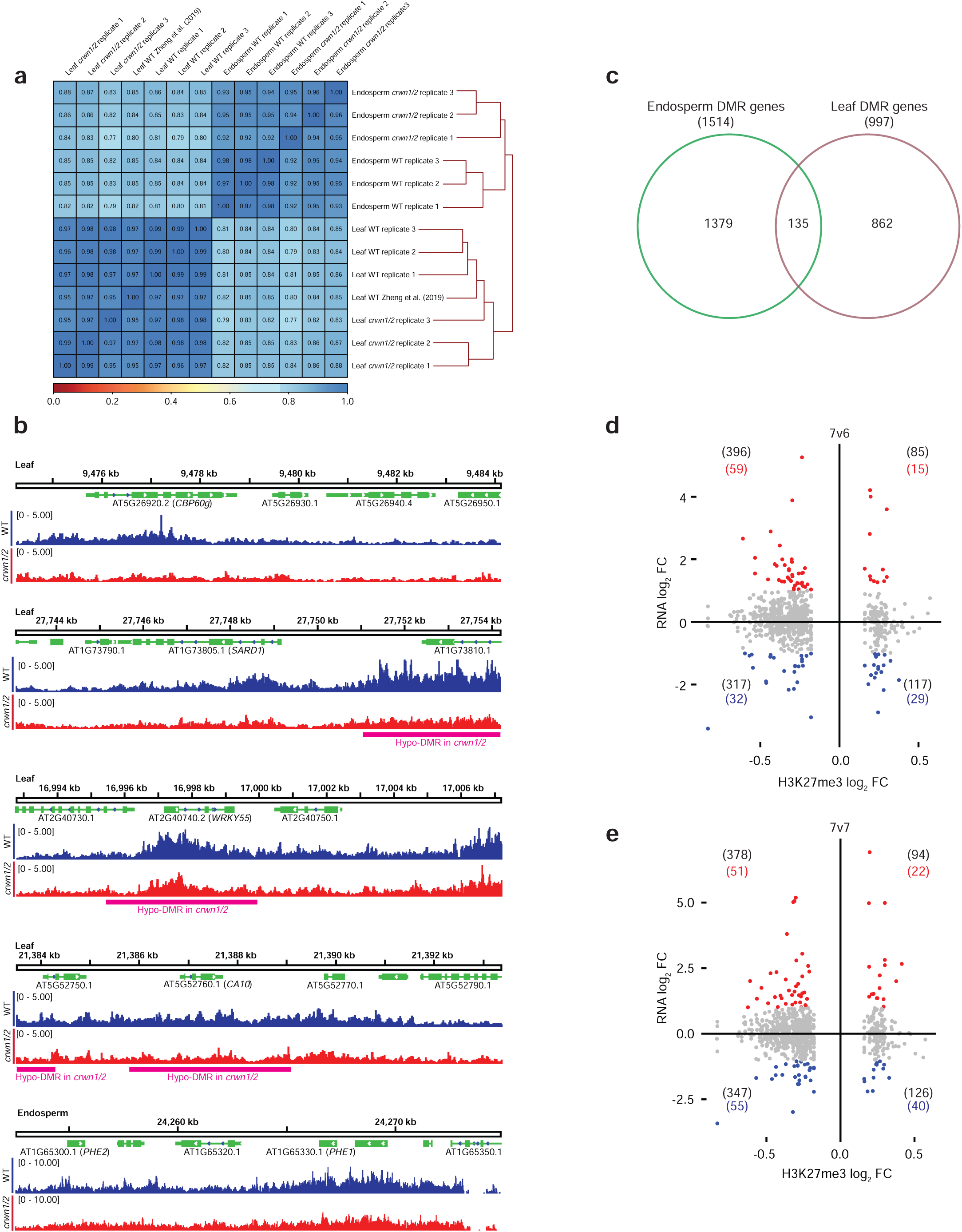
Correlation matrix of H3K27me3 CUT&RUN data, IGV browser snap shots of selected loci, and expression level of DMR genes. **a,** Pearson correlation matrix of H3K27me3/H3 signals across samples. Zheng and Gehring (2019) leaf H3K27me3 CUT&RUN data were included for comparison. **b,** IGV browser snap shots of leaf *CBP60g*, *SARD1*, *WRKY55*, *CA10,* and endosperm *PHE1*. DMRs defined by DiffBind are marked by magenta bars at the bottom of each snapshot. **c,** Venn diagram of endosperm and leaf DMR genes. **d, e,** The differential expression of DMR genes as a log_2_ fold change (FC) in *crwn1/2* compared to WT (regardless of *adjusted p* value) in 7vs6 (**d**) and 7vs7 (**e**) RNA-seq comparisons are plotted against their log_2_ FC in H3K27me3 level. Numbers in parentheses are number of DMR genes in each quadrant (black, regardless of *adjusted p* value). RNA log_2_ FC >= 1 (red, regardless of *adjusted p* value), RNA log_2_ FC <= -1 (blue, regardless of *adjusted p* value).

**Extended Data Fig. 6.**
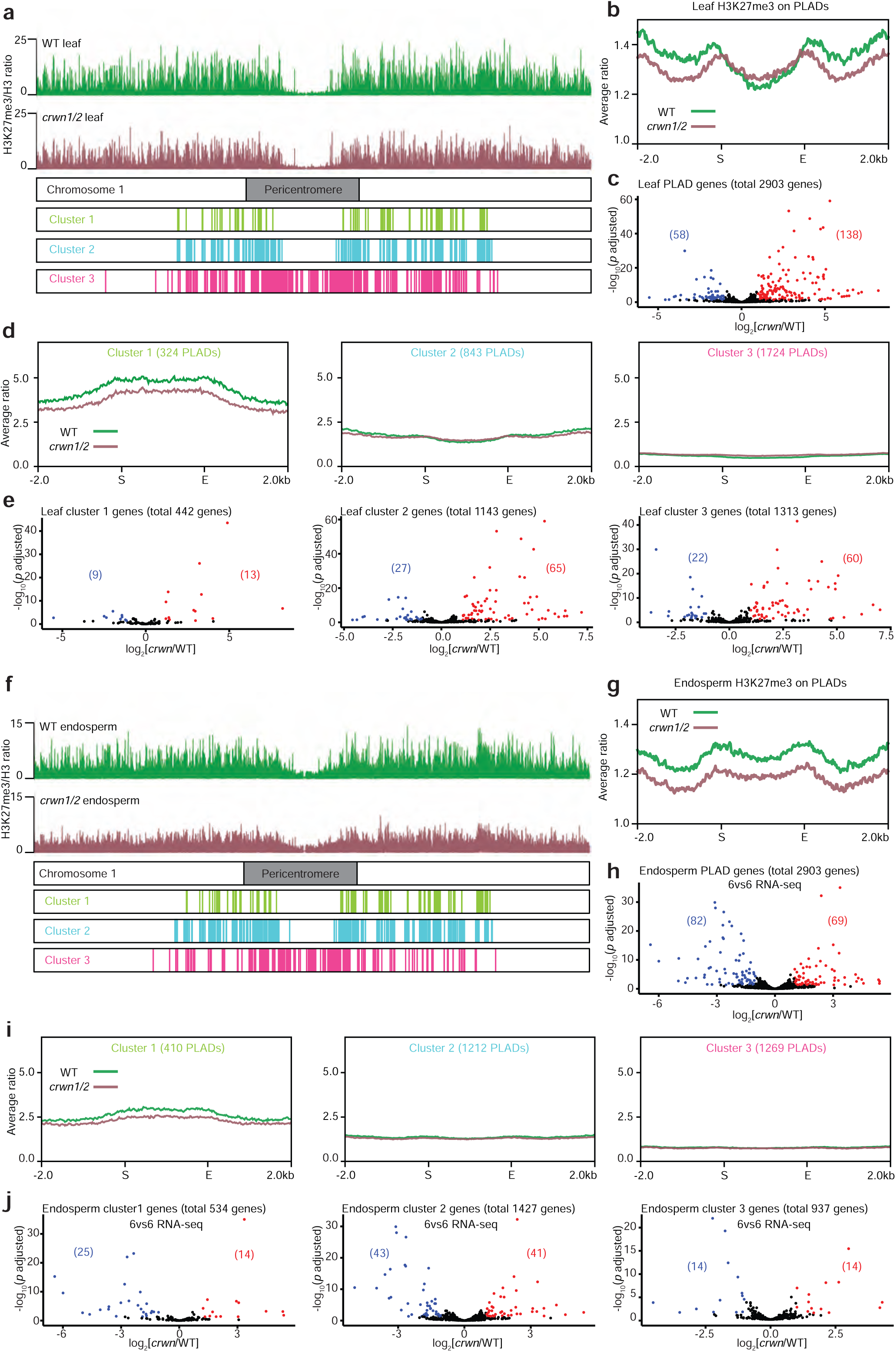
*k*-means clustering of PLADs based on their H3K27me3 and differentially expressed genes in each cluster. **a, f,** H3K27me3/H3 ratio in leaves (**a**) and endosperm (**f**) across chomosome 1 and PLAD clusters. H3K27me3/H3 signal ratio peaks (20bp bin size) are marked on top of chromosome diagrams. *k*-means clustering (*k* = 3) resolved PLADs into three groups based on their H3K27me3 level. Below the chromosome diagram, chromosomal loci of each PLAD cluster are marked. **b, g,** Average H3K27me3/H3 signal ratio on PLADs in leaves (**b**) and endosperm (**g**). **c, h,** Differentially expressed PLAD genes (|log_2_| >=1, *p adjusted* < 0.05) in *crwn1/2* leaves (**c**) and endosperm 6vs6 comparison (**h**). **d, i,** Differentially expressed PLAD genes (|log_2_| >=1, *p adjusted* < 0.05) in *crwn1/2* leaf PLAD clusters (**d**) and endosperm PLAD clusters (**i**). **e, j,** Volcano plots show differentially expressed PLAD genes in each leaf PLAD cluster (**e**) and endosperm PLAD cluster (**j**). PLAD genes are defined as those with at least 80% of their gene length (from 5’ UTR to 3’ UTR) overlapping with PLADs. Red, differentially expressed PLAD genes with increased expression (log_2_ >= 1, *p adjusted* < 0.05) in *crwn1/2* leaves (**e**) and endosperm (**j**) compared to WT. Blue, differentially expressed PLAD genes with decreased expression (log_2_ <= -1, *p adjusted* < 0.05) in *crwn1/2* leaves (**e**) and endosperm (**j**) compared to WT. Numbers in brackets indicate the number of genes in the same color category.

